# The ventral disc is a flexible microtubule organelle that depends on domed ultrastructure for functional attachment of *Giardia lamblia*

**DOI:** 10.1101/213421

**Authors:** Christopher Nosala, Scott C. Dawson

## Abstract

The parasite *Giardia lamblia* interacts with its host by directly attaching the lumen of the small intestine. Attachment is mediated by a cytoskeletal structure termed the ventral disc and proceeds in four distinct stages: skimming, seal formation, cell body contacts, and bare area contacts. The precise mechanism of disc-mediated attachment is unclear and attachment models rely heavily on whether or not the ventral disc is a dynamic structure. We sought to investigate the second stage of attachment in which a seal is formed beneath the ventral disc. Three-dimensional, live imaging of *Giardia* expressing specific ventral disc markers to the lateral crest, ventral groove, and disc body indicate dynamic movement in all of these regions. We observe seal formation by the lateral crest and determine that movement of the ventral groove region aids lateral crest seal formation. We also report the discovery of a new protein that is necessary for ventral disc formation and functional attachment (DAP_7268). Lastly, we observed that attachment largely depends on ventral disc ultrastructure as flattened discs display hindered attachment proficiency whether or not they retain the ability to form a seal. We propose a synthesized mechanism for attachment that includes flagellar hydrodynamic flow to help generate suction as well as disc conformational dynamics to aid in both hydrodynamic flow and the maintenance of negative pressure.

## Introduction

*Giardia lamblia* is a zoonotic intestinal parasite that causes significant diarrheal disease worldwide (1). *Giardia*sis disproportionately impacts people in developing countries where early and recurrent childhood infection is associated with worsened malnutrition and delayed development. Trophozoites are the motile flagellated form of the parasite that readily attach to the lumen of the small intestine. During colonization, *Giardia* forms a monolayer but does not invade cells nor tissues. The trophozoite’s interphase cytoskeleton comprises many microtubule structures including the ventral disc, four pairs of bisymmetrical flagella, the median body, and the funis (2). The ventral disc mediates attachment within the host gut and is essential for *in vivo* colonization because unattached trophozoites are swept away via peristalsis and shed in the feces where they cannot survive harsh environmental conditions (3). Recent studies indicate that *Giardia* can attach to a variety of surfaces regardless of surface treatment (PEG, Teflon) (4). This indifference to surface chemistry may help explain *Giardia*’s zoonotic potential by allowing the parasite to attach to a variety of mammalian intestinal environments despite varying luminal chemistry. It remains unclear exactly how *Giardia* performs attachment, yet this mechanism allows quick reversible adherence to almost any surface.

The foundation of the ventral disc is a parallel array of microtubules ~100 polymers thick (Figure 1A). Built on these microtubules are distinct substructures including outer microtubule associated proteins, inner microtubule associated proteins, microribbons, and crossbridges (5). Although both outer and inner MAPs are discernible via cryo-ET, the identity of these MAPs remains unknown (6, 7). The microribbons are trilaminar sheets of protein that jet dorsally into the cell body from the microtubules. Crossbridges join the microribbons laterally with regular spacing. These ventral disc substructures are found throughout the entirety of the ventral disc yet the role they play in forming ventral disc ultrastructure and functional attachment, as well as the protein makeup of each substructure, is largely unknown. Eighty five proteins localize to the ventral disc (5). How these proteins are involved in building and maintaining the ventral disc structure is unclear. Many GFP localizations correlate with differential protein densities described using cryo-tomography (6, 7). These observations indicate the ventral disc can be separated into distinct regions: disc body, overlap zone, ventral groove, disc margin, and the lateral crest (Figure 1A).

**Figure 1:**
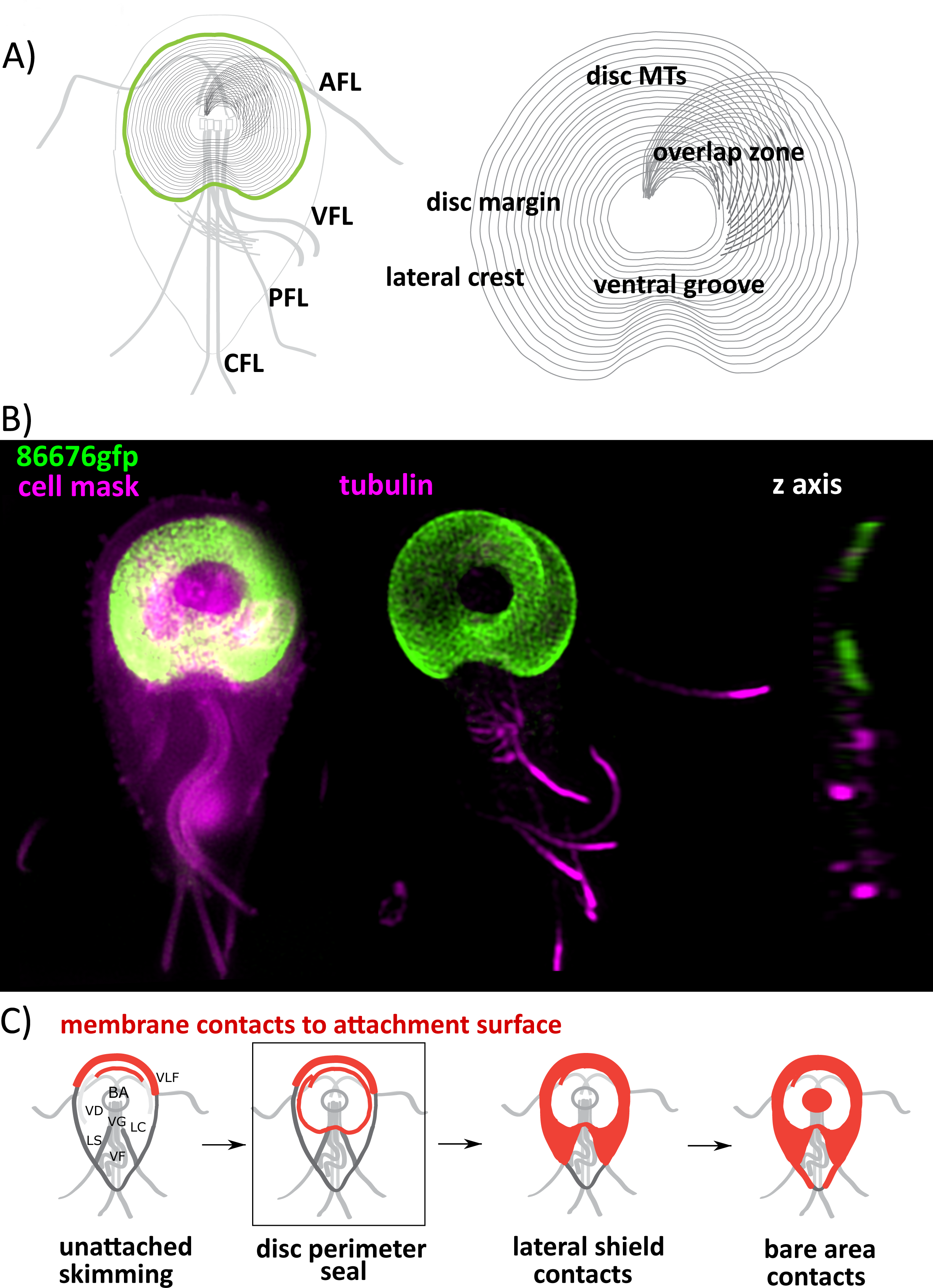
The ventral disc comprises distinct regions and is responsible for progressing through the stages of attachment. *Giardia* cells contain four pairs of flagella as well as the ventral disc. Ventral disc microtubules spiral away from the flagellar basal bodies through distinct regions: the disc body, the ventral groove, the disc margin, and the overlap zone. The lateral crest surrounds the outside of the disc (1a). 3D-SIM microscopy of a GFP-tagged protein known to localize to the disc (DAP_86676, delta-giardin) indicates the disc’s position within the cell and the disc’s domed shape (1b). Attachment progresses through four distinct stages defined by membrane contacts with the attachment surface: ventrolateral flange (VLF), lateral crest (LC), bare area (BA), ventral groove (VG), ventral flagella (VF), ventral disc (VD), lateral shield (LS). Skimming cells display VLF contacts before a complete seal is formed beneath the LC and VD. Next, LS contacts increase as the cell body approaches the surface and finally the BA contacts the surface.

Ventral disc mediated suction is the prevailing attachment model because *Giardia* can attach regardless of surface chemical treatment, because attachment is rapidly reversible, and because trophozoites prefer flat to bumpy surfaces (anything else?). Suction would require a means of generating negative pressure beneath the domed shape of the disk as well as the formation and maintenance of a seal. TIRF microscopy has been used to view *Giardia*’s membrane interaction with the attachment surface and *Giardia* was observed to progress through four distinct stages of attachment (Figure 1C) (8). These attachment stages proceed in reverse during detachment. Early in attachment, trophozoites encounter the attachment plane and cells orient ventral disc side down. Parasites then skim along the surface with membrane/surface contact at the anterior portion of the ventrolateral flange. The cell membrane is observed to form a seal when a suitable habitation site is encountered. This seal progresses from the anterior part of the cell to surround the entire ventral disc area with the portion near the ventral groove contacting the surface last. There remains many unanswered questions regarding this stage of attachment. Is the ventral disc an active player in seal formation? Is seal formation necessary for attachment? Why is the ventral groove region the last to contact the surface? What can specific ventral disc regions teach us about *Giardia*’s attachment mechanism?

We sought to investigate the second stage of attachment in which the membrane forms a contiguous seal beneath the ventral disc. Prior descriptions of ventral disc function rely heavily on static images of fixed cells (EM) or indirect observations of membrane dynamics (5). We used a variety of live imaging strategies with *Giardia* expressing specific ventral disc markers to clarify a long standing controversy regarding ventral disc conformational changes. We observe seal formation by the lateral crest as well as dynamic movement of the ventral groove region *in vivo*. Here we report the discovery of a new protein that is necessary for ventral disc formation and functional attachment (DAP_7268). Lastly, we observed that attachment largely depends on ventral disc ultrastructure as flattened discs display hindered attachment proficiency whether or not they retain the ability to form a seal.

## Results

### Flagella are important for establishing but not maintaining attachment

We developed a novel shear stress assay using a commercial flow chamber (Ibidi mSlide VI 0.4) to quantify *Giardia* attachment forces (Figure 2A, movies Supplemental 1). In this assay we used the 86676-gfp strain because the fluorescent ventral disc signal allowed for rapid quantification of cell number before and after shear force was applied. 86676-gfp cells were challenged using a variety of forces and it was found that 90% of these cells were able to resist 2ml/min of flow (~4dyn/cm2).

**Figure 2:**
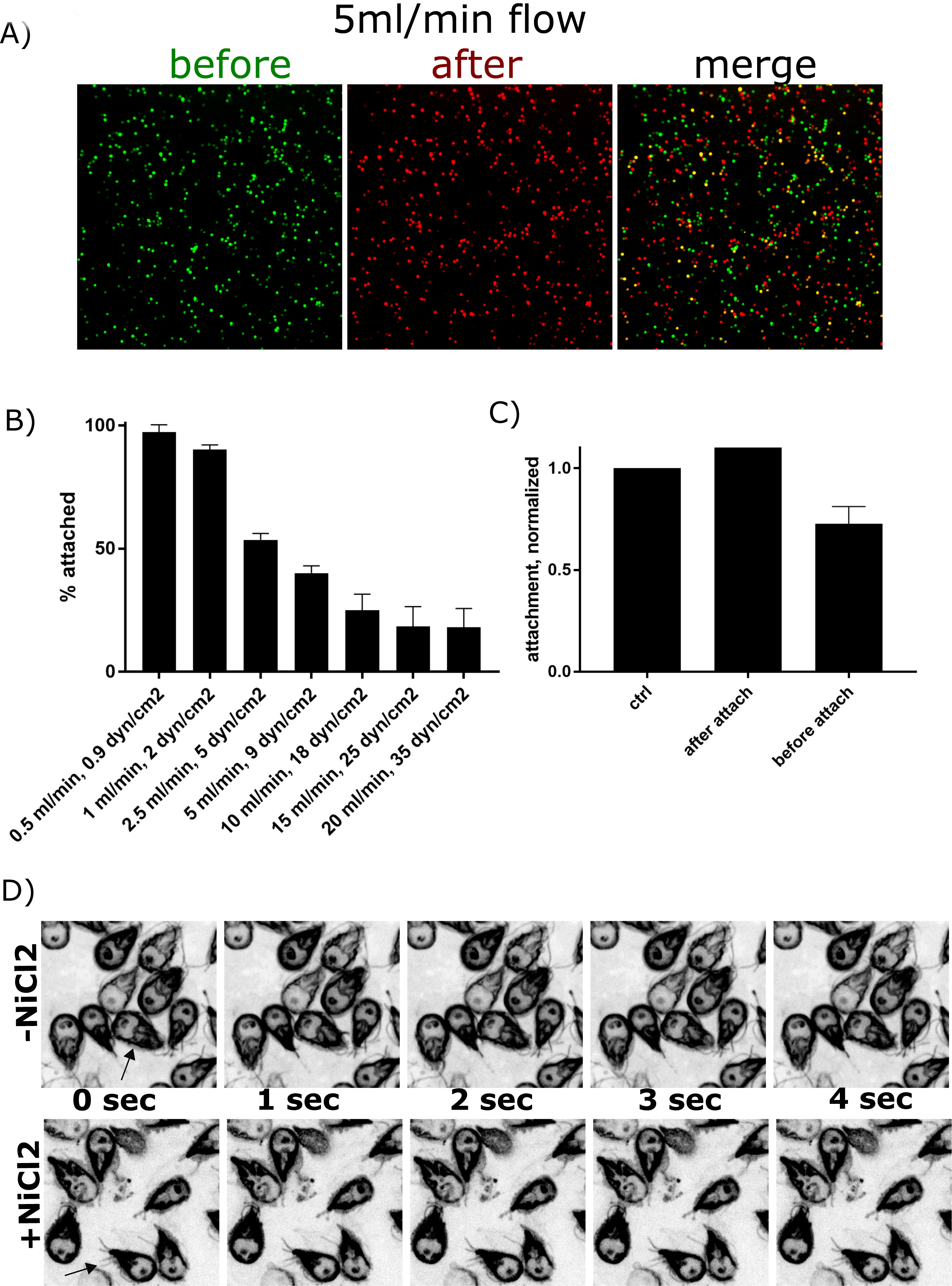
Flagellar beating contributes to initiating but not maintaining attachment. We developed a novel flow assay to challenge attached cells with shear stress. Fluorescent cells (86676gfp) are allowed to attach prior to being challenged with a specific flow rate. Before and after pictures are collected and false colored green and red, respectively (2a). Percent of cells that can resist the shear stress challenge are calculated and 90% of cells were able to resist 2ml/min flowrate (2b). NiCl_2_ treatment effectively ceases flagellar beating (2d). Cells were treated with NiCl_2_ after attaching in the flow chamber or before attachment. Cells treated before attachment showed a significant decrease ability to resist 2ml/min flow (T-test, p≤0.05).

Using two genetic techniques to disrupt normal flagellar beating, our lab has previously reported that the ventral flagella are not required to maintain attachment of *Giardia* trophozoites to surfaces (8). We used a pharmacological approach to confirm these findings and to test if the ventral flagella contribute to initiating attachment. NiCl2 has been used in a variety of systems to inhibit flagellar beating (9). Treatment of *Giardia* trophozoites with NiCl2 inhibits beating of all eight flagella (Figure 2D). Attached trophozoites treated with NiCl2 retain proper membrane surface contacts including a complete seal and bare area.

Control cells were allowed to attach for 30 mins prior to challenge with 2 ml/min flow and all tests were normalized to control attachment efficiency (Figure 2C). To test if the flagella were important for maintaining attachment, cells were allowed to attach for 30 mins, 25mM NiCl2 was then added to the chamber, and attached cells were challenged with 2 ml/min flow once flagellar beating ceased. No significant difference was observed in attachment deficiency consistent with the observation that ventral flagellar beating is not required for maintaining attachment. To test if the flagella were important for initiating attachment, cells were pre-incubated with 25mM NiCl2 prior to addition to the flow chamber and allowed to attach for 30 mins before challenge with 2 ml/min flow. We observed a significant 27% reduction in attachment efficiency for pre-treated cells compared to control cells indicating ventral flagellar beating is important for initiating attachment.

### The lateral crest imparts a seal to fluid flow

In an ongoing GFP screen, our lab has discovered forty proteins that localize to either the outer disc margin or the lateral crest. These proteins display variability in their localization that emphasizes both the ventral groove and overlap regions (5). We hypothesize that the lateral crest surrounding the outside of the ventral disc is responsible for seal formation. This hypothesis is supported by previous DAP_16343 knockdown data wherein open discs with broken lateral crests do not properly progress through the stages of attachment and display erroneous membrane/surface contacts (8).

We utilized diffusion of fluorescent microspheres (FluoSpheres) to investigate fluid flow and ventral disc seal formation in attached cells (Figure 3). Trophozoites were allowed to attach and FluoSpheres were added just prior to imaging. Time lapse videos were collected (Figure 3A) and projected to view diffusion of FluoSpheres around the attached cells (Figure 3C). Consistent with seal formation, beads rarely entered into the space directly below the ventral disc. Beads that did enter underneath the disc did so exclusively at the ventral groove region indicating fluid flow at this region (Figure 3A). After entering the space below the disc, beads could ricochet off the lateral crest perimeter and exit at the ventral groove region parallel with the ventral flagella. These observations are consistent with a hydrodynamic model in which the ventral flagella draw fluid out from underneath the ventral disc to generate negative pressure.

**Figure 3:**
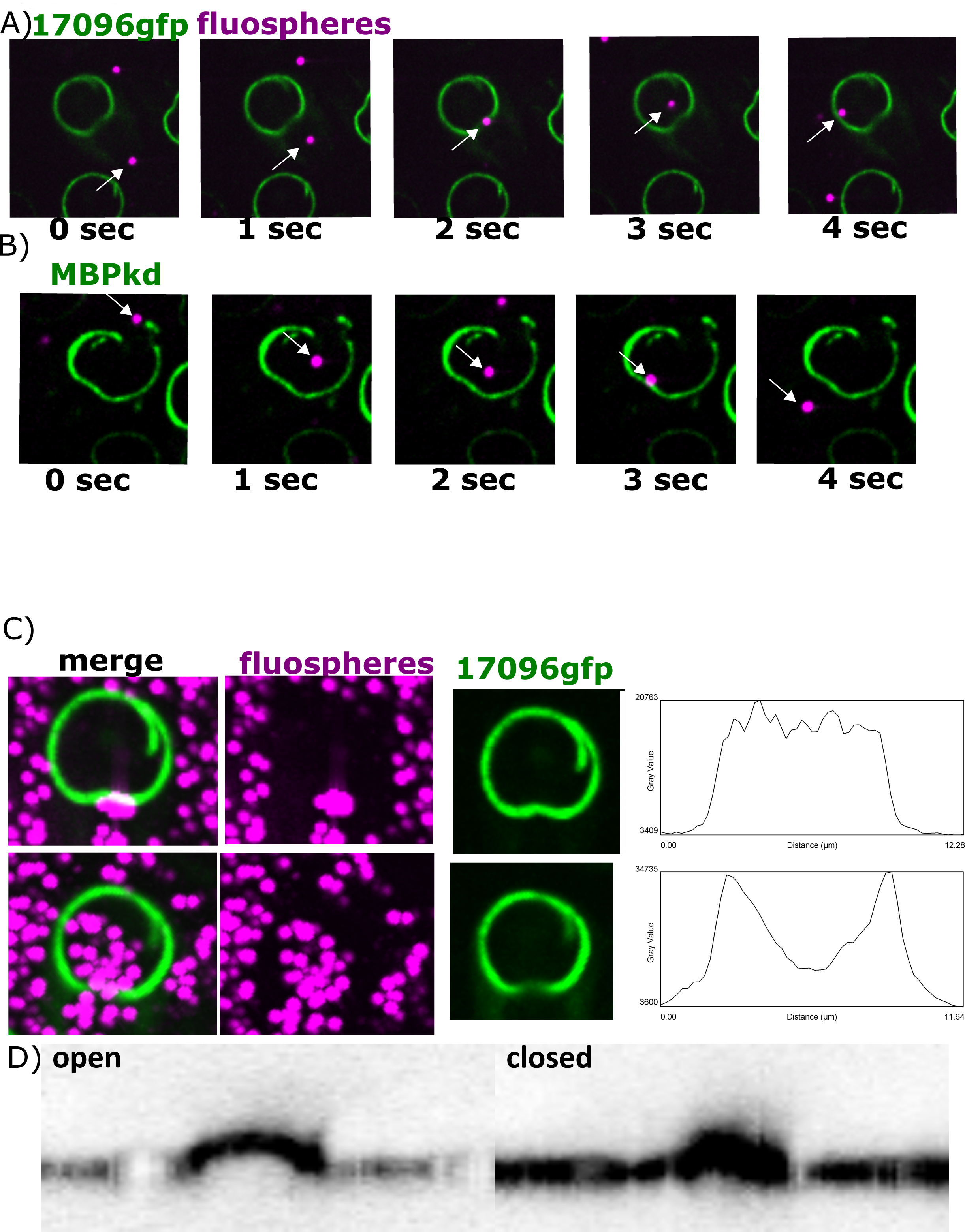
Fluid flow in and out of the compartment below the ventral disc is governed by the disc margin. Cells expressing an outer disc marker (DAP_17096-gfp) were allowed to attach before addition of fluorescent microspheres (FluoSpheres) and diffusion of FluoSpheres was recorded. In some cases, miscrospheres entered the compartment beneath the disc at the ventral groove indicating fluid flow at this region (3a). In cases of discontinuity (MBPkd), beads were pulled into the disc compartment and swept out of the ventral groove parallel with the ventral flagella (3b). FluoSphere diffusion was recorded by imaging every second for five minutes. A maximum intensity projection of this timecourse indicates total FluoSphere diffusion (3c). Line scans at the ventral groove indicate a decrease in signal at the ventral groove in cases where FluoSpheres entered the disc compartment. Three-dimensional imaging of the ventral groove indicates both open and closed states (3d).

We repeated the bead diffusion assay in and DAP_16343 knockdown cells and observed a severe defect in the ability of only DAP_16343 cells to form a seal (Figure 3B). FluoSpheres were observed to freely enter areas of lateral crest breakage in DAP_16343 knockdown cells. After entering underneath the disc, FluoSpheres were swept out from under the disc in line with the ventral flagella (Figure 3B). This observation is consistent with the idea that the ventral flagella serve to pull fluid out from underneath the disc in order to generate negative pressure.

Line scans indicate a decrease in GFP signal at the ventral groove region for cells under which beads entered, consistent with an ‘opened gate’ state (Figure 3C). In cases where beads approached the ventral groove region but did not enter, GFP signal was consistent with a closed ventral groove state. We observed the same result using three different candidate markers of the lateral crest (13981-gfp, 17096-gfp, 17231-gfp) (supplemental data). Three dimensional imaging of recently attached trophozoites indicates both open and closed states at the ventral groove region (Figure 3D).

### Dynamic movement of ventral groove region

We next wanted to investigate the lateral crest during attachment of trophozoites to determine whether the lateral crest is a driver of seal formation. Kymographs of time lapse videos at the plane of attachment using three different markers (13981-gfp, 12139-gfp, 86676-gfp) indicate increases and decreases of GFP signal specifically at the ventral groove region consistent with movement in the Z-axis (Figure 4). Kymographs of detaching cells indicate that the ventral groove region rises to break the surface seal prior to other areas of the lateral crest and lastly the entire cell detaches. We performed the same experiment using the 86676-gfp strain to determine if the observed movement was specific to the outside of the disc or the entire disc body. The ventral disc was observed to change between flattened and domed shapes and the most dramatic movement was observed at the ventral groove region (Figure 4C).

**Figure 4:**
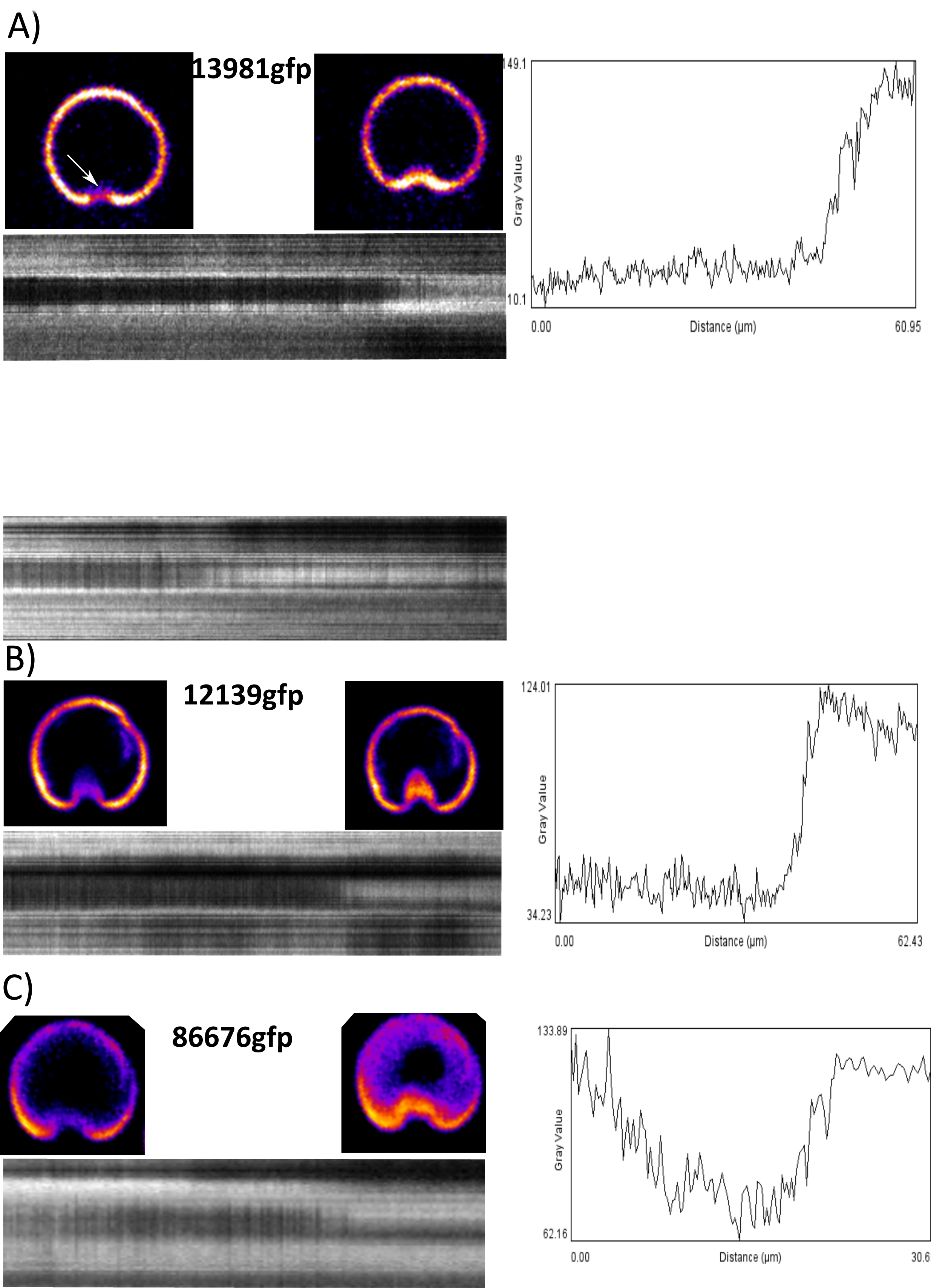
Dynamic movement is observed at the ventral groove region. Three different ventral disc markers display movement at the ventral groove region: 13981gfp (lateral crest), 12139gfp (disc margin, ventral groove), 86676gfp (disc body). Movies were recorded by imaging every second for five minutes. Kymographs were generated by tracing a line around the outer edge of the disc through the ventral groove region (black dotted line) and reslicing through the entire timecourse. Line plots were generated by drawing a line through the ventral groove region of the kymograph (white dotted line).

### DAP_7268 is necessary for domed ventral disc ultrastructure but not lateral crest formation

Morpholino anti-sense oligo based knockdown is the current standard for gene knockdown in *Giardia* (8, 10, 11). This tool has been used previously to disrupt the ventral disc architecture using anti-DAP_16343 morpholino (Figure 5a) (12). We found that morpholino knockdown of a new ventral disc protein, DAP_7268, results in a disrupted disc phenotype that differs from DAP_16343 knockdown (Figure 5a). In the DAP_86676-gfp (dGiardin) background strain, knockdown of DAP_7268 results in a break in the left lobe of the ventral disc body that is visible in the Z axis (Figure 5C). This break appears to be regarded as a ventral disc edge because the outer disc proteins DAP_13981-gfp and DAP_12139-gfp localize to this region (Figure 5A, Supp). Unlike the open disc phenotype typical of DAP_16343 knockdown cells, DAP_7268 knockdown cells retain a fully contiguous lateral crest (Figure 5A). Fluorescent beads were not observed to enter underneath the disc of DAP_7268 knockdown cells supporting the idea that this phenotype retains a fully functional seal.

**Figure 5:**
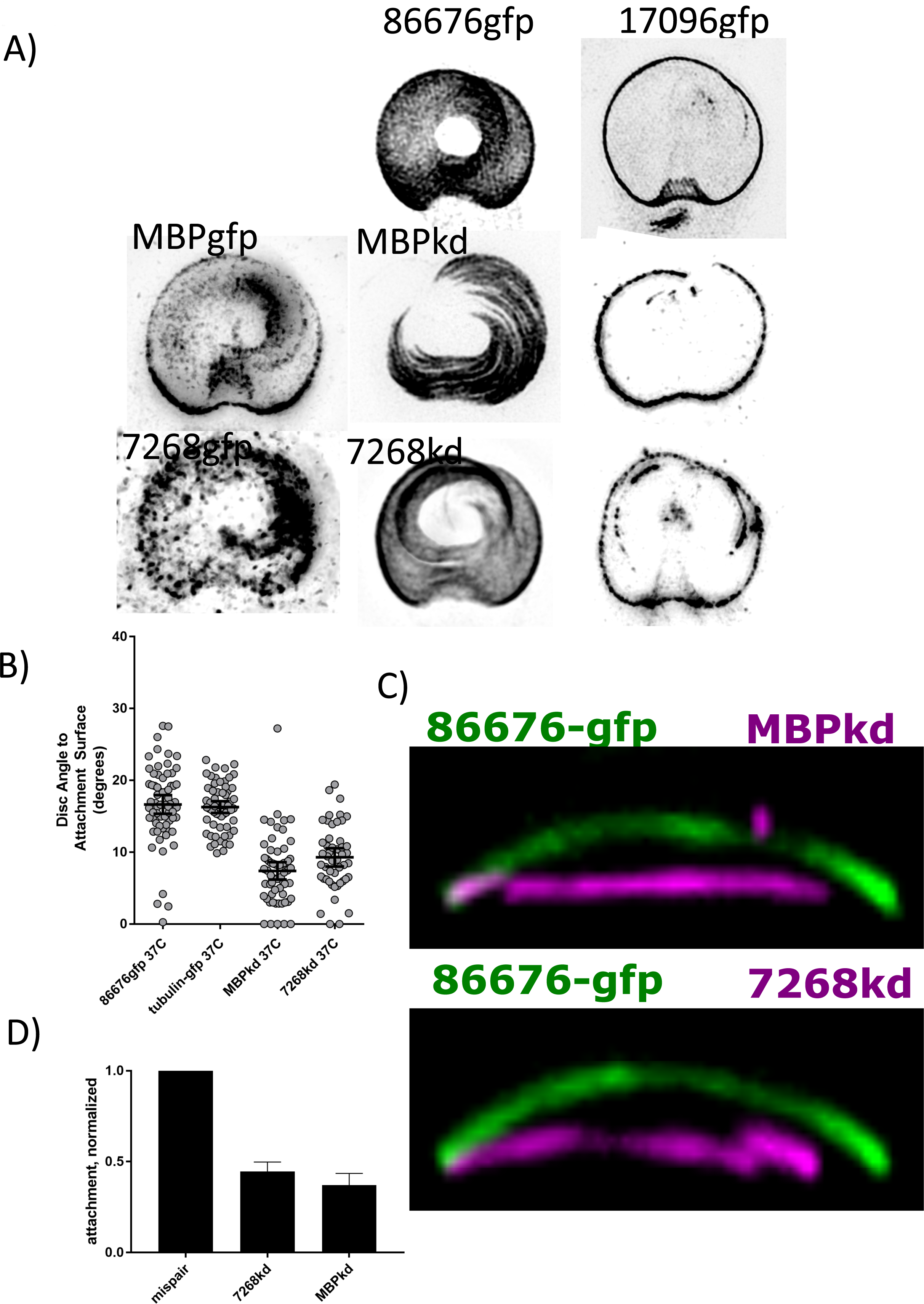
Knockdown of DAP_16343 and DAP_7268 result in flattened discs that poorly resist shear stress challenge. DAP_86676gfp and DAP_17096gfp were used as markers of the disc body and lateral crest, respectively. Knockdown of DAP_16343 (MBP) results in incompletely formed discs with a broken lateral crest. Knockdown of DAP_7268 results in a broken disc that retains a completely formed lateral crest, as well as extraneous lateral crest deposition (5a). Disc curvature was quantified in three dimensional images by measuring the angle of the disc relative to the attachment surface in the DAP_86676gfp and beta-tubulin-gfp strains (~18deg). Knockdown of both DAP_16343 and DAP_7268 results in flattening of the ventral disc to ~8deg (5B,C). Flow challenge of knockdown cells at 2ml/min was normalized to mispair morpholino control and indicates severe defects in ability to resist shear stress (5D).

The domed shape of the ventral disc has been hypothesized to be important for functional attachment (12). We quantified ventral disc ultrastructure by measuring the angle at which the ventral disc contacts the attachment surface. Two different ventral disc markers (dGiardin-gfp, aTubulin-gfp) indicated an average angle of ~17degrees ventral disc angle for attached cells at 37C. Both DAP_7268 and DAP_16343 knockdown cells were significantly flattened to ~8degrees compared to parent DAP_86676gfp cells (Figure 5B,C).

### Knockdown phenotypes poorly resist shear stress

We next wanted to ask whether the ability to form a lateral crest seal offers an attachment benefit to DAP_7268 knockdown cells compared to DAP_16343 knockdown cells in the shear stress flow assay. DAP_16343 knockdown results in a flattened disc and incompletely formed lateral crest seal whereas DAP_7268 knockdown results in a flattened disc that retains a fully formed lateral crest seal (Figure 5a). Both DAP_16343 knockdown and 7268 knockdown cells displayed significant defects in their ability to resist 2ml/min of flow in our assay. We found that the ability to form a seal did not offer an attachment advantage to 7268 knockdown cells as there was no significant difference in the ability of 7268 knockdown cells to resist shear flow compared to DAP_16343 knockdown cells (Figure5d).

### Ventral disc curvature is critical for attachment

We sought to determine whether ventral disc curvature could explain the attachment defects observed in our knockdown cells. *Giardia* trophozoites grown in laboratory conditions are commonly removed from culture vessels by placing the tubes on ice until cells detach. To reproduce this phenomenon, imaging plates were placed on ice and immediately imaged to observe the curvature of the disc within ‘detached’ cells. We observed a significant decrease in the ventral disc angle after incubation on ice to ~8degrees indicating the ventral disc’s domed ultrastructure is flexible and flattened in iced detached cells (Figure 6A).

**Figure 6:**
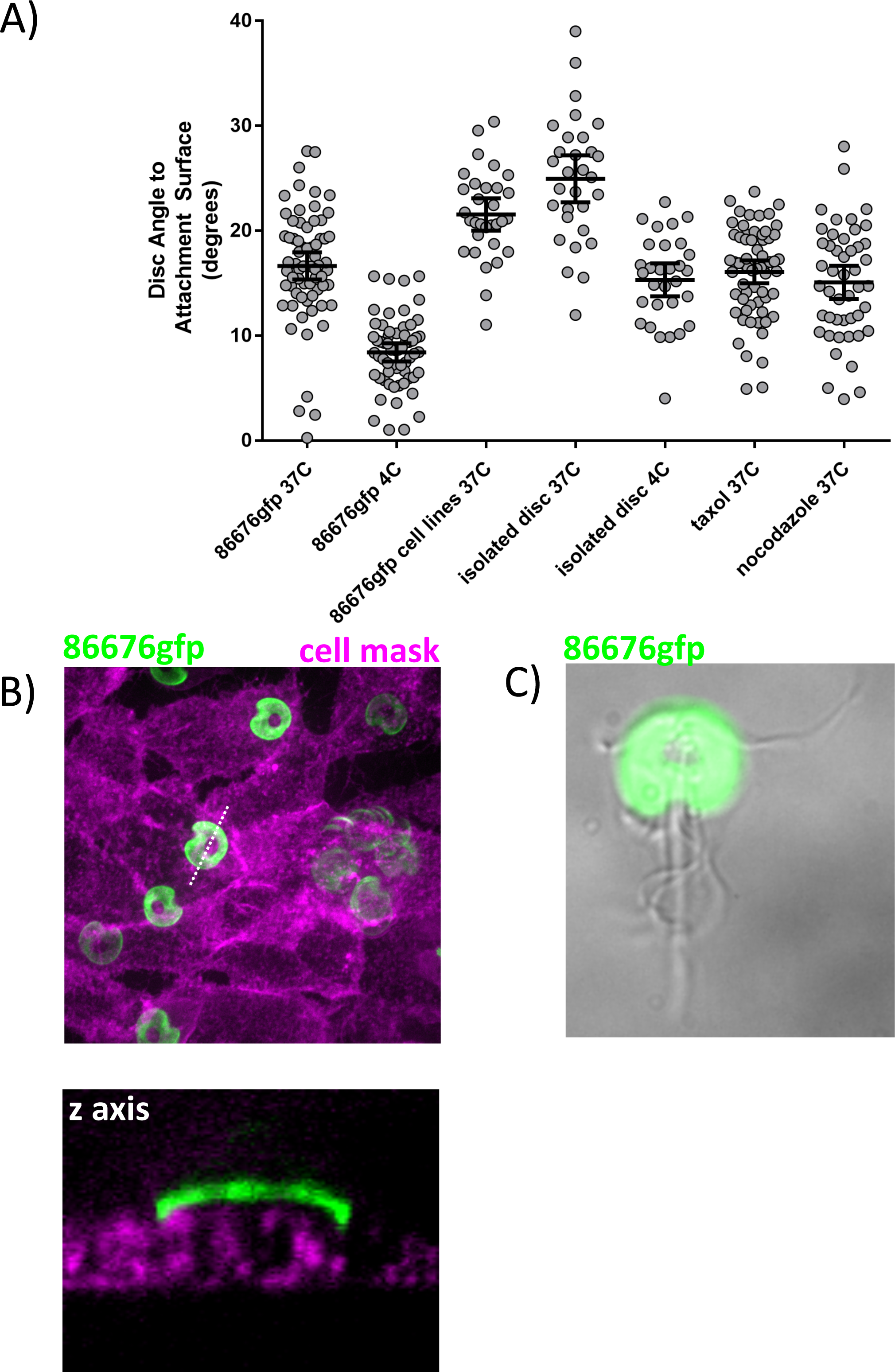
Disc curvature is sensitive to temperature changes. DAP_86676gfp was used as a ventral disc marker to measure ventral disc curvature. Cells subjected to cold temperatures (4C) result in flattened disc that mimic knockdown angles (~8deg). Cells attached to human cell lines display an increase in ventral disc curvature (~22deg) and detergent-extracted ventral discs display an even greater increase in curvature (~25deg). Isolated ventral discs subjected to cold temperature displayed a decrease in curvature from ~25deg to ~18deg. Drugs targeting microtubule dynamic instability (taxol, nocodazole) had no significant effect on disc curvature in live cells.

**Figure 7:**
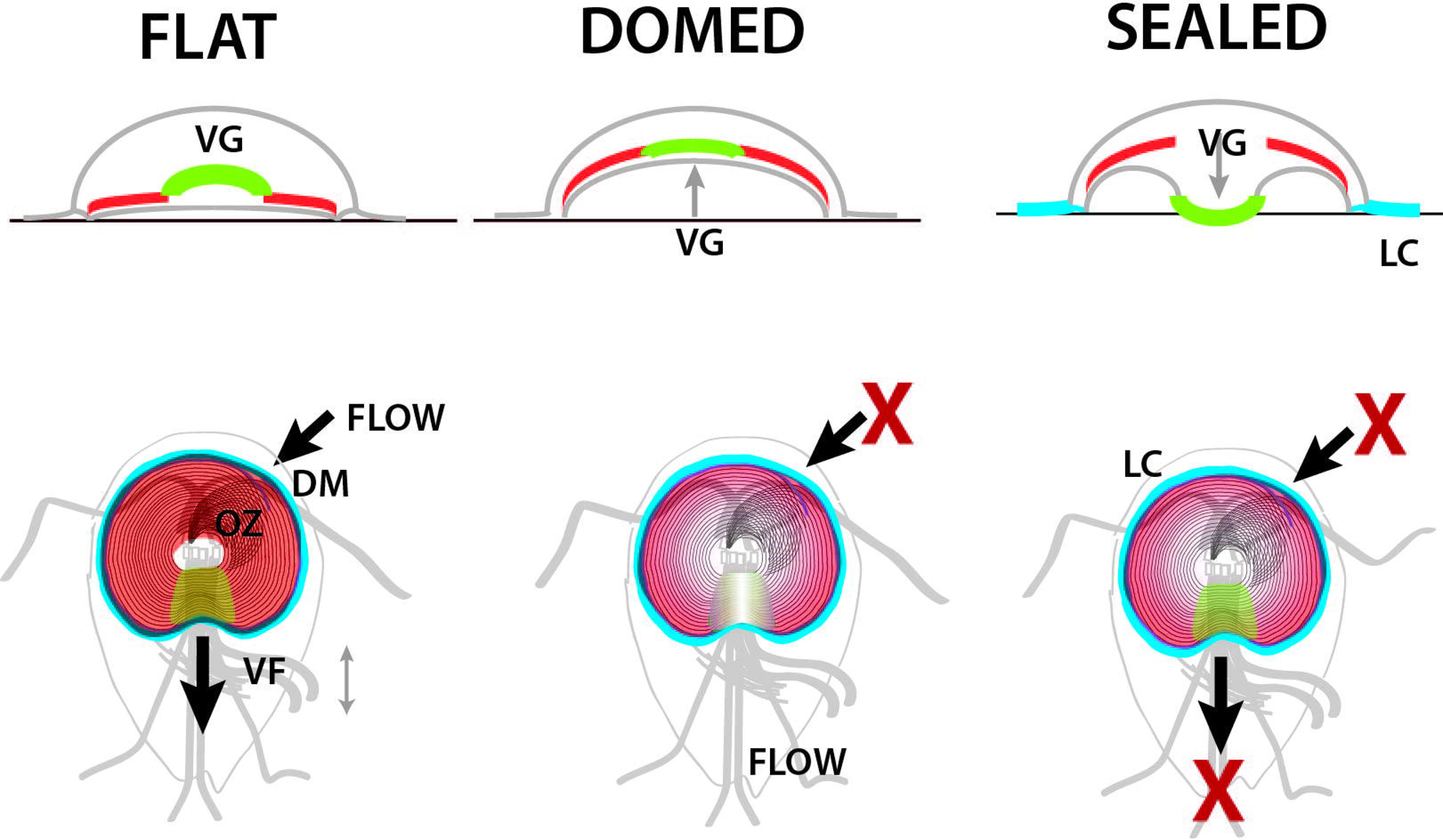
Disc conformational model of *Giardia* attachment. Conformational changes of the ventral disc, together with ventral flagellar beating, generate the forces of hydrodynamic suction that enable attachment. Ventral flagellar beating established hydrodynamic flow (arrows). In early attachment, the disc adopts a flattened conformation that opens the disc margin and ventral groove regions (FLAT), allow flow underneath the disc. Disc doming then creates space underneath the disc, simultaneously closing the disc margin and raising the ventral groove (DOMED), modulating hydrodynamic flow. Lastly, the ventral groove closes prior to the formation of a disc perimeter seal by the lateral crest (SEALED), limiting flow underneath the disc.

The rigidity of the attachment surface may play a role in the degree to which ventral discs generate curvature. We cultured *Giardia* on a monolayer of MCF 10A human epithelial cells (ATCC CRL-10317) to determine the ventral disc attachment angle on human cell membranes that are more deformable than glass or plastic. Ventral discs displayed a small yet significant increase in curvature when attached to the human cells. Consistent with suction, human cell membranes appeared to be pulled into the space underneath the disc (Figure 6B). We next wanted to determine whether the ventral disc structure is curved per se or if curvature is dependent on cellular factors. *Giardia*’s microtubule cytoskeleton, including the ventral disc and all eight flagella, can be readily extracted using detergent buffers (Figure 6C). Isolated ventral discs displayed a significant increase in curvature to ~25 degrees at 37C. We imaged these isolated ventral discs after incubating on ice to determine if temperature alone is sufficient to drive disc conformational changes. Cold isolated ventral discs displayed a significant decrease in curvature compared to warm isolated ventral discs from 17degrees to 8degrees, although not to the same degree as seen within live cells indicating that cell factors or physical constraints within the cell likely contribute to ventral disc curvature *in vivo*. (Microtubule dynamic instability has not been observed within the ventral disc, although small changes to microtubule polymerization could help explain disc conformational dynamics. Treatment of warm attached cells with the drugs taxol and nocodazole that target microtubule dynamic instability did not have an effect on ventral disc attachment angle.)

## Discussion

Of the 85 proteins that are currently known to localize to the ventral disc (5), only two have been found to be essential for disc biogenesis and/or attachment. DAP_16343 (MBP, median body protein) was first described to be essential and served as a proof of principle that knockdown of a single DAP could disrupt ventral disc ultrastructure and affect attachment (12). Exactly why knockdown of 16343 results in an incompletely formed disc with collapsed microtubule/microribbon spacing remains to be determined. Little is known about the timing of ventral disc biogenesis, how ventral disc microtubule and microribbon polymerization is controlled, how proteins are targeted to specific regions, etc. Regulation of ventral disc biogenesis is particularly interesting considering two new ventral discs are generated in a relatively short time during cell division (10). Differing knockdown phenotypes can help us investigate both ventral disc biogenesis and attachment mechanisms.

Here is the first description that DAP_7268 is an essential ventral disc protein. Knockdown of DAP_7268 resulted in a less drastic ventral disc structural phenotype than knockdown of DAP_16343, although both exhibited similar attachment defects in our shear stress test. The break in the ventral disc occurred in the same relative position of the disc body for every 7268 knockdown cell observed. This crack may be mistaken as a ventral disc edge by the DAP_7268 knockdown cells because proteins that localize to the lateral crest/disc margin also localize to this region (Figure 5A). Ankyrin structural motifs are predicted within 33 ventral disc protein (13), including DAP_7268, which has ankyrin repeats at both the N and C termini (5). Ankyrin repeats are believed to be facilitators of protein/protein interactions so DAP_7268 could be a structural component of a ventral disc substructure. Future studies will determine which ventral disc substructure each DAP localizes to in order to assign function to these substructures in disc biogenesis and functional attachment.

The ventral disc is a microtubule structure that is unique to *Giardia* species, although some DAPs have homologs in *Giardia*’s closest relative Spironucleus spp. Tubulin is conserved across eukaryotes. *Giardia* aTubulin shares <u>94%</u> sequence identity with human tubulin. How has *Giardia* adapted conserved components to build the ventral disc, a completely novel organelle with novel function? Some hypothesize that the ventral disc is a modified flagellum, consistent with ventral disc microtubules sharing the basal bodies with flagellar microtubules. However, most of the proteins contained within the ventral disc are categorized as ‘hypothetical’ because they do not share homology with any known characterized proteins (5). It is likely these novel proteins have allowed for the invention of this novel organelle. Hypothetical proteins in the ventral disc are important from a disease perspective because proteins that are necessary for ventral disc biogenesis and functional attachment are not found in the human genome making them good candidate drug targets.

The ventral disc structure was first viewed in detail using EM by Cheissin in 1964 (14). This marks the beginning of the controversy over whether or not the disc is a dynamic structure as Cheissin opines that the ventral disc structure appears considerably elastic. The ventral disc of *Giardia* muris attached to the mouse intestine was next observed by Friend in 1966 (15). Friend argues that the disc appears very rigid and that the disc itself is not responsible for attachment, a point we disagree with based on our knockdown phenotypes and live imaging. In a series of papers beginning 1973, Holberton used EM and live imaging to observe the ventral disc and the waveform of the ventral flagella of *Giardia* muris (16), and develops the hydrodynamic model of *Giardia* attachment that has prevailed for decades. In this model, lateral channels are posited to exist outside the disc and the beating of the ventral flagella causes fluid flow through these channels to the ventro-caudal groove to generate hydrodynamic attachment forces. However, the lateral channels necessary for hydrodynamic fluid flow have not been observed in live cells. Our lab previously sought to test the hydrodynamic model using two genetic methods that disrupt normal ventral flagellar beating (8). In that study, ventral flagellar beating was observed not to be necessary to maintain attachment of *Giardia* to surfaces, although the flagella were observed to be important for initially establishing attachment and orienting the cell to the attachment plane. Consistent with these earlier results, FluoSphere flow in our bead diffusion assay did not mimic fluid flow through possible lateral channels.

It is necessary to consider that *Giardia muris* and *Giardia lamblia* are different species with different host specificities and may rely on different, albeit similar, attachment mechanisms. The hydrodynamic model relies on a constantly open ventral groove region that is lacking in *Giardia muris* (see old EM, (16), EM of both species in 1981 (17). Many proteins in *Giardia lamblia* specifically target to the ventral groove region, and these proteins are lacking in the *Giardia muris* genome (personal correspondence, Staffan Svard). GFP localization patterns suggest that there are distinct substructures in the ventral groove region, each composed of unique proteins.

Despite positing that the ventral disc of *Giardia muris* is a rigid structure (16, 18, 19), Holberton later argued that the ventral groove region (posterior notch) is flexible in isolated discs of *Giardia lamblia* (17). Holberton observed flexible unwinding of this region with and without the addition of ATP, arguing that this flexibility is not active but has stored elastic energy (17). Our time lapse imaging of attaching *Giardia lamblia* trophozoites shows clear movement of the ventral groove *in vivo* that is consistent with flexibility of this region that we believe is important for the forming and breaking of a seal during attachment. It remains to be determined how movement of the ventral groove region is facilitated *in vivo*.

Our observations indicate that temperature is sufficient for altering ventral disc curvature and that cold discs are flattened out. This data helps explain why placing *Giardia* culture vessels on ice results in detachment of the parasite. The temperature/disc shape relationship may ensure parasites only attach within living hosts. Isolated ventral discs displayed an increase in curvature indicating that a domed shape is not actively generated but rather is inherently built into the ventral disc structure. It also remains to be determined if ventral discs actively flatten out during routine attachment. Recent advances in optical imaging (Lattice Light Sheet) will allow for direct imaging of ventral disc dynamics in real time.

If suction is the main mechanism of attachment, and the lateral crest is responsible for forming a seal, then why do some MBP knockdown cells with severe lateral crest defects remain attached? In these cases the outer membrane shows a clear blebbing that fills in the gap. This membrane could serve as a redundant seal that allows for suction in the event of lateral crest disruption.

Ventral disc attachment mechanism that we argue for is akin to that described by Cheissin in 1964 in which the incessant beating of the ventral flagella pulls fluid out from under the disc (14). Together with a flexible ventral disc and mobile lateral crest, a seal is formed that secures the parasite to the attachment surface by the negative pressure that is generated. Therefore, the lateral crest acts as a barrier that does not allow fluid to enter or exit underneath the disc, except when a break is generated at the ventral groove. This ventral groove is the region at which we observed movement of FluoSpheres to enter and exit the space underneath the ventral disc. The opening and closing of this gate is also consistent with our observations that the ventral flagella contribute to establishing attachment, but are not necessary for maintaining attachment.

## Materials and Methods

### *Giardia* Culture

*Giardia* intestinalis strain WBC6 (ATCC 50803) trophozoites were maintained at 37°C in modified TYI-S_33 medium with bovine bile (20)in 16-ml screw-cap tubes (Fisher Scientific). Upon reaching confluency, the strain was split by first placing tubes on ice for 15 minutes then transferring 0.5ml of detached culture to 11.5ml of warmed media. Prior to imaging, cells were washed 3x with warm 1xHBS to remove autofluorescence associated with culture medium.

### *Giardia* Strain Generation

All strains were constructed as previously described (21). For C-terminal GFP episomal tag: All candidate DAP PCR forward primers (see Table S1) were designed to bind 200 bp upstream of the gene to include the *Giardia* native promoter and contained the sequence CACC at the 5’ end to facilitate directional cloning. Blunt-ended PCR amplicons were generated by PCR using PfuTurbo Hotstart PCR Mastermix (Stratagene) with *Giardia* intestinalis strain WBC6 genomic DNA. The candidate DAP PCR amplicons were subsequently subcloned into the Invitrogen pENTR/D-TOPO backbone to generate Gateway entry clones. Inserts in entry clones were sequenced to confirm the identity and correct orientation of the gene. To construct DAP-GFP fusions, positive entry clones were then recombined, via LR reaction, with a 1-fragment GFP tagging E. coli/*Giardia* shuttle destination vector (pcGFP1F.pac) using LR Clonase II Plus (Invitrogen). LR reactions were performed using 100 ng pcGFP1F.pac and 150 ng of DAP entry clone plasmid DNA. Positive clones were screened by digestion with AscI, and bulk plasmid DNA was prepared using Qiagen’s Plasmid Midi Kit. To create C-terminal GFP-tagged candidate DAP strains, *Giardia* intestinalis strain WBC6 was electroporated with roughly 20 mg of plasmid DNA (above) using the GenePulserXL (BioRad) under previously described conditions. Episomal DAP-GFP constructs were maintained in transformants using antibiotic selection (50 mg/ml puromycin).

### Morpholino Gene Knockdown

Morpholinos were designed to target the first 24 bases of the gene and 1 base upstream of the 5’ end. Morpholino sequences used in this study are as follows antiDAP_16343: GCTGAAAACCATAGCCTCGGACATT, anti7268: GAACCAGTCGCTCGCGGTTTGCATG. Prior to morpholino knockdown, trophozoites of DAP86676-GFP, DAP17096-GFP, and DAP12139-GFP-tagged strains were grown to confluence. Cells were washed with 12 ml of fresh medium, quickly suspended again in 12 ml of medium, and then chilled on ice for 15 mins. The detached trophozoites were then centrifuged at 900xg for 5 min at 4°C. The resulting pellet was resuspended in 0.27 ml of fresh medium. This was added to a 0.4-cm electroporation cuvette along with 0.03 ml of morpholino at a final concentration of 100mM. The cuvette was chilled on ice for 15 min, electroporation was performed, and the cells were placed back on ice for 15 mins. These transformed cells were then incubated at 37°C for 24 h. Cell morphology and attachment were assessed in both live and fixed trophozoites after morpholino knockdown.

### Structured Illumination Microscopy

3D stacks were collected at 0.125 um intervals on the Nikon N-SIM Structured Illumination Super-resolution Microscope with a 100x/NA 1.49 objective, 100 EX V-R diffraction grating, and an Andor iXon3 DU-897E EMCCD. Images were recollected and reconstructed in the “2D-SIM” mode (no out of focus light removal; reconstruction used three diffraction grating angles each with three translations).

### Confocal Microscopy

3D stacks and time lapse movies were acquired of live cells grown in 96-well #1.5 black glass bottom imaging plates (In Vitro Scientific). Images were acquired with the spinning-disk module of a Marianas SDC Real-Time 3D Confocal-TIRF microscope (Intelligent Imaging Innovations) fit with a Yokogawa spinning-disk head, a 63×/1.3 NA oil-immersion objective, and electron-multiplying charge-coupled device camera. Acquisition was controlled by SlideBook 6 software (3i Incorporated). All raw images were exposed and scaled with the same parameters.

### Ventral Disc Angle Measuements

Cells were allowed to attach to imaging plates at 37C for 30 mins in 1x HBS, washed 3x with 1xHBS to remove unattached cells, and 100 ul of warm 1xHBS was added. Next, 100ul of 37C 3% low melt agarose in 1xHBS was added to a final concentration of 1.5% agarose to immobilize cells on the imaging plates. Immobilization was not observed to affect disc curvature. Slices were taken at 0.2 um intervals for 8 um to capture the entire disc. Disc curvature images was generated by reslicing the stack laterally across the posterior portion of the bare area and angles were measured in FIJI. At least 30 cells were measured on three separate days totaling 90 cells for each condition.

### FluoSphere Diffusion Assay

Cells were allowed to attach to imaging plates at 37C for 30 mins in 1x HBS, washed 3x with 1xHBS to remove unattached cells, and 200 ul of warm 1xHBS was added. FluoSpheres Carboxylate-Modified Microspheres (0.2 um, red fluorescent) beads were pretreated for 24 hours by transferring 5 ul from stock to 1 ml 1xHBS containing 1%BSA and stored at 4C to help prevent clumping before use. At time of assay, 20 ul of this diluted solution was added to each well and time lapse imaging began immediately. Single focal planes were acquired every 1 s for 5 minutes with definite focus. Image analysis was performed in FIJI.

### Shear Force Assay

Metamorph image acquisition software (MDS Technologies) was used to collect single focal plane images using a Leica DMI 6000 wide-field inverted fluorescence microscope with a 10x objective. Fluorescent images were collected before and after challenge with flow for shear stress. DIC time lapse movies were collected during flow challenge by imaging the same focal plane every second for 60 seconds. A fresh chamber was used for each assay in Ibidi mSlide VI 0.4 flow chambers. Cells in frame were imaged for 20 seconds, challenged with flow for 20 seconds, followed by 20 seconds more video before final fluorescent image was collected. Attached cells were counted by overlaying the before image (false colored green) over the after image (false colored red). Green cells were counted as unable to resist flow challenge whereas yellow cells were able to resist challenge. Percent attached was calculated by (#yellow)/(#yellow + #green). In some cases, percentages were normalized to control from a parallel run.

## Acknowledgements

Human MCF 10A cells were a kind gift from Nont Kosaisawe and John Albeck (Dept. MCB, UC Davis). Thank you to Michael Paddy of the UC Davis MCB Microscopy Core with helpful advice on SDC and SIM microscopes. Kari Hagen, Shane McInally, and Eric Walters provided helpful readings of manuscript drafts.

## References

1. Einarsson E, Ma'ayeh S, & Svard SG (2016) An up-date on Giardia and giardiasis. Current opinion in microbiology 34:47–52.

2. Dawson SC (2010) An insider's guide to the microtubule cytoskeleton of *Giardia*. Cellular microbiology 12(5):588–598.

3. Nosala C & Dawson SC (2015) The Critical Role of the Cytoskeleton in the Pathogenesis of Giardia. Curr Clin Microbiol Rep 2(4):155–162.

4. Hansen WR, Tulyathan O, Dawson SC, Cande WZ, & Fletcher DA (2006) *Giardia lamblia* attachment force is insensitive to surface treatments. Eukaryot Cell 5(4):781–783.

5. Nosala CaD, S.C. (2017) “Disc-o-Fever”: getting down with Giardia’s groovy microtubule organelle. Trends In Cell Biology in press.

6. Brown JR, Schwartz CL, Heumann JM, Dawson SC, & Hoenger A (2016) A detailed look at the cytoskeletal architecture of the Giardia lamblia ventral disc. J Struct Biol 194(1):38–48.

7. Schwartz CL, Heumann JM, Dawson SC, & Hoenger A (2012) A detailed, hierarchical study of *Giardia lamblia*’s ventral disc reveals novel microtubule-associated protein complexes. PLoS One 7(9):e43783.

8. House SA, Richter DJ, Pham JK, & Dawson SC (2011) Giardia flagellar motility is not directly required to maintain attachment to surfaces. PLoS Pathog 7(8):e1002167.

9. Ginger ML, Portman N, & McKean PG (2008) Swimming with protists: perception, motility and flagellum assembly. Nature reviews. Microbiology 6(11):838–850.

10. Hardin WR, et al. (2017) Myosin-independent cytokinesis in Giardia utilizes flagella to coordinate force generation and direct membrane trafficking. Proc Natl Acad Sci U S A 114(29):E5854–E5863.

11. Carpenter ML & Cande WZ (2009) Using morpholinos for gene knockdown in Giardia intestinalis. Eukaryot cell 8(6):916–919.

12. Woessner DJ & Dawson SC (2012) The Giardia median body protein is a ventral disc protein that is critical for maintaining a domed disc conformation during attachment. Eukaryot Cell 11(3):292–301.

13. Li J, Mahajan A, & Tsai MD (2006) Ankyrin repeat: a unique motif mediating protein-protein interactions. Biochemistry 45(51):15168–15178.

14. Cheissin EM (1964) Ultrastructure of Lamblia Duodenalis. I. Body Surface, Sucking Disc and Median Bodies. J Protozool 11:91–98.

15. Friend DS (1966) The fine structure of Giardia muris. J Cell Biol 29(2):317–332.

16. Holberton DV (1973) Mechanism of attachment of Giardia to the wall of the small intestine. Transactions of the Royal Society of Tropical Medicine and Hygiene 67(1):29–30.

17. Holberton DV & Ward AP (1981) Isolation of the cytoskeleton from *Giardia*. Tubulin and a low-molecular-weight protein associated with microribbon structures. Journal of cell science 47:139–166.

18. Holberton DV (1974) Attachment of *Giardia*-a hydrodynamic model based on flagellar activity. J Exp Biol 60(1):207–221.

19. Holberton DV (1973) Fine structure of the ventral disk apparatus and the mechanism of attachment in the flagellate Giardia muris. Journal of cell science 13(1):11–41.

20. Keister DB (1983) Axenic culture of *Giardia lamblia* in TYI-S-33 medium supplemented with bile. Trans. R. Soc. Trop. Med Hyg. 77:487–488.

21. Hagen KD, et al. (2011) Novel structural components of the ventral disc and lateral crest in Giardia intestinalis. PLoS Negl Trop Dis 5(12):e1442.

